# Origin of co-operativity in the activation of dimeric transcription factors

**DOI:** 10.1101/769661

**Authors:** Martin Welch, Jens Christian Brasen, Christopher T. Workman, Thomas Sams

## Abstract

Cooperative behavior in the binding of ligands to a protein is often viewed as a complex phenomenon where conformational changes induced by the binding of the first ligand leads to tighter binding of subsequent ligands. We revisit the ligand-dependent activation of dimeric transcription factors and show that this process may appear cooperative even when it results from independent ligand binding events. This effect is further accentuated through binding of the activated transcription factor to its cognate operator site on the DNA, where we demonstrate that cooperative activation is a stable fixed point. Our analysis nicely accounts for the apparent co-operativity inherent in the biological activity of many dimeric transcription factors.

## I. INTRODUCTION

The “central dogma” of Biology describes how genetic information flows from DNA to RNA, and thence to protein [1]. DNA information encoded by genes is therefore “expressed” via two tightly regulated processes: transcription, where the static DNA sequence is converted into a transient RNA “message”; and translation, where the information carried by the RNA messenger is decoded to yield functionally-active protein. The process of transcription is largely controlled by “transcription factors” [2]. Transcription factors are capable of recognizing discrete DNA sequence motifs in regulatory regions called “operators” located just upstream of the gene(s) of interest. Once bound to these operator regions, the transcription factor either recruits (if it is acting to stimulate expression of the gene) or blocks (if it is acting to repress expression of the gene) access of the RNA-synthesizing enzyme, RNA polymerase, to its binding site on the DNA. The binding site of RNA polymerase on a gene is known as the “promoter”, and this is usually closely juxtaposed with the operator. This way, binding of a transcription factor to an operator site can alter the transcription rate of the associated gene. If the gene encodes a protein, the elevated rate of transcription yields more messenger RNA, which, in turn, yields proportionally more protein following translation.

Because of its duplex form, DNA has intrinsic structural dyad symmetry. This means that many transcription factors function as dimers, with each monomer within the dimer recognizing sequence elements on just one strand of the duplex. Furthermore, the activity of many transcription factors is dependent upon the binding of low molecular weight “co-inducers”. These small molecules (ligands) are thought to bind to the transcription factor and alter its conformation or oligomeric state, thereby altering the affinity of the transcription factor for the DNA. One may therefore express the activity of the transcription factor by the occupancy of the operator as a function of ligand concentration. Indeed, many transcription factors are now characterized into discrete classes based on the type of ligand bound by the first member of each family to be historically identified. We therefore encounter “LysR-type” or “LuxR-type” regulators, for example.

Below, we will revisit the process of forming active transcription factor dimers as a consequence of ligand binding to the protein and subsequent binding of the activated transcription factor-ligand complex to the DNA. Our analysis was inspired by a bacterial cell-cell communication mechanism called “quorum sensing”. In quorum sensing, a freely-diffusible self-produced signal molecule (frequently, an *N*-acylated homoserine lactone (AHL)) accumulates in the culture. Once the AHL concentration exceeds a critical threshold concentration (thought to be determined primarily by binding of the AHL to a LuxR-type transcription factor), the transcription factor becomes activated and elicits the transcription of key target genes, which are often involved in bacterial virulence [3, 4]. The resulting phenotypic “switch” is often very abrupt, and has all the hallmarks of being underpinned by a highly cooperative mechanism [5–8]. However, there is currently no obvious mechanism by which such cooperativity might be achieved – a problem that we aim to address directly in the current study. Furthermore, our findings can be readily extended to other classes of transcription factor, and may well therefore be general.

The modeling of quorum sensing systems has been challenged by the controversy as to whether the LuxR-type transcription factors associated with quorum sensing are of the common form where transcription factor dimerization drives ligand binding, or whether some are of the form where ligand binding drives transcription factor dimerization [9–12]. The controversy was partially resolved when Sappington and co-workers [13] demonstrated that, *in vivo*, LasR folds reversibly into its dimer form prior to reversibly binding ligands. This is consistent with the kinetic study reported by Claussen *et al.* [14]. We shall therefore adopt (as a starting point) the situation where dimerization of the transcription factor precedes ligand binding(s). A formalism for the case where ligand binding precedes dimerization is dealt with in the appendix.

While looking into the details of the transcription factor activation, we noted that a system driven by continuous production and turnover of the transcription factor leads to a rescaled dissociation constant and increased cooperativity. In the current work, we derive an expression for the effective dissociation constant of the ligand-transcription factor complex, and establish a proper independence measure for the ligand bindings. The formalism clarifies how it is possible to discriminate between cooperative and non-cooperative activation, and between fast and slow response to changes in ligand concentration, depending on the rate of transcription factor turnover.

Perhaps the earliest quantitative explanation for ligand binding to a protein was proposed by Hill in his analysis of the oxygen transport properties of haemoglobin. Here, the form 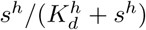, where *s* is the ligand concentration, *K*_*d*_ is the dissociation constant, and *h* is the “cooperativity coefficient” (also known as the Hill coefficient) provides a good description of receptor occupancy at ligand concentrations close to the *K*_*d*_ [15–17]. However, the Hill expression fails to describe the full dynamic range of an activated transcription factor. In the current work, we remedy this by showing that the concentration of the activated dimer with two ligands bound, *r*_4_=[R_2_S_2_], can be approximated by the expression;

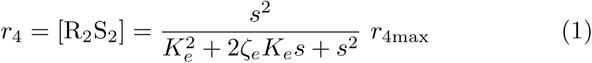

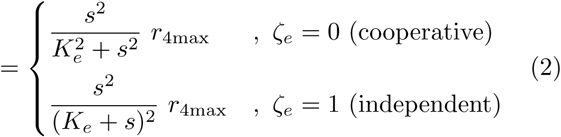

where *s* = [S] is the ligand concentration, *K*_*e*_ denotes the effective dissociation constant (which may differ from the underlying dissociation constant for ligand binding to the dimeric transcription factor), and where *ζ*_*e*_ is the independence measure, which lies between 0 (cooperative ligand bindings) and 1 (a product of two independent ligand bindings). The maximal concentration for the active transcription factor, *r*_4max_, is reached when ligand concentration is much larger than the effective dissociation constant.

## II. ANALYSIS

Many transcription factors function as homo- or heterodimers. We shall primarily be concerned with the typical case where homodimerization drives ligand binding. However, for completeness, a brief account of the case where ligand binding precedes dimerization is presented in the appendix.

### A. Dimerization drives ligand binding

The kinetics describing constitutive production of the transcription factor (R), the dimerization of the transcription factor, and its activation trough ligand (S) binding are;

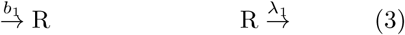

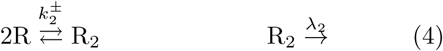

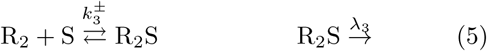

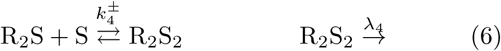

as illustrated in Fig. 1. The first step in the process is the constitutive production of transcription factors at rate *b*_1_. This step encompasses transcription from the DNA segment into messenger RNA as well as translation of the messenger RNA into transcription factors. These enzymatic processes are carried out by the RNA polymerase and the ribosomes respectively. In Fig. 1 the gene segment that codes for the transcription factor is indicated as a framed *R* and its associated promoter *P*_*R*_. The net result of this is the constant production of transcription factors at rate *b*_1_ as indicated in Eq. (3). Eqs. (4)-(6) describe the dimerization of the transcription factor and activation through ligand bindings into the active form, R_2_S_2_, of the transcription factor complex. They are all reversible reactions between diffusible molecules. On-rate constants are superscripted “+” and off-rates are superscripted “−”. In Fig. 1 green arrows indicate the forward reactions (i. e. the “+”). In the first reaction, two transcription factors associate with rate constant 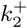 labeling at both green arrows leading to the formation of R_2_. Similarly for the two ligand bindings. The activated transcription factor complex, i. e. R_2_S_2_, is able to bind to its cognate operator region and control the transcription of a gene (*X* in Fig. 1). In Fig. 1 this regulation is indicated by the binding of the activated transcription factor to the operator (*P*_*X*_) in the regulatory region of the gene on the DNA. We will return to this final step in section II C. The rightmost processes in Eqs. (3)-(6) are the turnover of the protein at rates *λ*_*i*_ which include dilution by cell division as well as protein degradation by the cell’s recycling system handled by specialised proteins (proteases). Note that we allow for dissociation of the dimerized transcription factor (R_2_). Should the dimer be stable, i. e. when 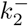 is negligible, the formalism still applies.

**FIG. 1.**
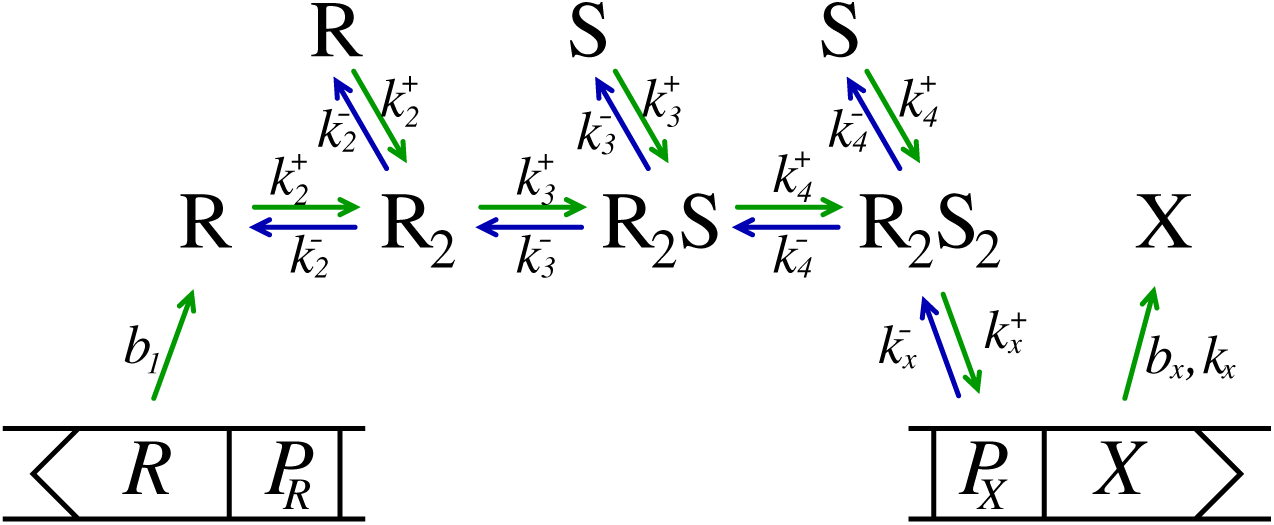
Activation of transcription factor by ligand binding. Schematic diagram of the production, dimerization, and activation of a dimeric transcription factor followed by the binding of the active form of the transcription factor, R_2_S_2_, to an operator site on the DNA controlling the production of substance X. The timeline for forward reactions reads from left to right and following green arrows. Blue arrows signify the time direction for reverse reactions when appropriate. The subscript number for the rate constants gives the overall time order of reactions corresponding to Eqs. (3)-(6). Association rate constants are superscripted “+” whereas dissociation rates are superscripted “−”. The monomeric transcription factor is denoted R while ligands are denoted S. The different forms of the transcription factor, i. e. R, R_2_, R_2_S, and R_2_S_2_ diffuse freely as does the ligands. The framed *R* signifies a piece of DNA that codes for the protein R, while the framed *P*_*R*_ is the promoter region on the DNA controlling the transcription of the gene. We now take a quick tour through the pathway. 1: The transcription factor, R, encoded by the gene *R* is produced at rate *b*_1_ (Eq. (3)). This involves two unidirectional processes, first transcription to messenger RNA then translation in to the transcription factor. 2: Two copies of the transcription factor bind to each other with rate constant 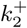 to form the dimer R_2_ (Eq. (4)). 3: The ligand and the transcription factor-dimer associates with rate constant 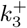 to form R_2_S (Eq. (5)). 4: Another copy of the ligand reacts with the R_2_S complex to form the activated transcription factor, R_2_S_2_, with rate constant 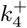 (Eq. (6)). 5: Finally, the activated transcription factor binds to the operator region controlling the promoter for transcription of the gene *X*. This shifts the production from background level, *b*_*x*_, to induced level, *k*_*x*_. The overall role of the regulatory motif is to control the expression of the product X in response to changing concentrations of the ligand S. At low concentrations of active transcription factors, the transcription of *X* proceeds at background rate, *b*_*x*_, and shifts to *k*_*x*_ at high concentrations of active transcription factors. In the manuscript, the concentrations (not shown) of constituents are *r*_1_=[R], *r*_2_=[R_2_], *r*_3_=[R_2_S], and *r*_4_=[R_2_S_2_]. The total concentration of promoter sites on the DNA is denoted *p*_*t*_=[*P*_*X*_] + [R_2_S_2_-*P*_*X*_] while the concentration of activated sites is *p*_*a*_=[R_2_S_2_-*P*_*X*_]. All forms of the transcription factors may be subject to proteolytic degradation (not shown).

If we denote the concentrations *r*_1_=[R], *r*_2_=[R_2_], *r*_3_= [R_2_S], *r*_4_=[R_2_S_2_], and *s*=[S], the kinetic equations corresponding to Eqs. (3)-(6) are;

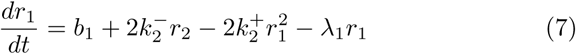

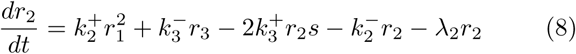

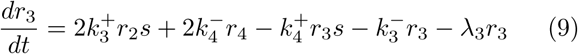

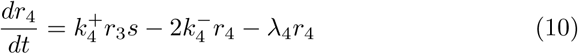

The explicit inclusion of the proper statistical weights (factors of 2) allows for simpler interpretation of independence in the analysis.

The maximal activated concentration, *r*_4max_, the effective dissociation constant, *K*_*e*_, and the independence measure, *ζ*_*e*_, at steady state, will now be determined. To determine the effective dissociation constant (*K*_*e*_) in the parametrization of the activated transcription factor in Eq.(1), we consider its asymptotic behavior at small and large values of *s* for the second order process. On a logarithmic scale, these are recognized as straight lines (of slope 2 and 0 respectively) and the effective dissociation constant sits at their crossing. For independent ligand bindings, *r*_4_ deviates by a factor of 4 from its maximal value at the crossing of the asymptotes, while it only deviates by a factor of 2 for cooperative ligand bindings. Its actual deviation at the crossing of the asymptotes will be used to determine the independence measure. The remainder of this section, concerns the mathematical details of this procedure.

When *s* is large, *r*_2_ and *r*_3_ are negligible and *r*_1_ assumes its minimal value;

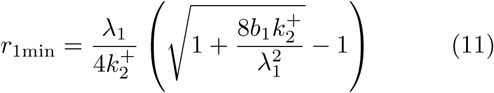

which can be seen by setting *dr*_1_*/dt* = 0 with *r*_2_ = 0. The maximal value of *r*_4_ is then;

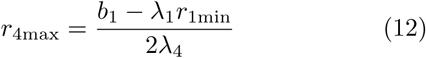

which is just a net dimer source term diluted by growth and degradation (*λ*_4_). When *s* is small *r*_1_ and *r*_2_ will be at their maximal value, and *r*_4_ will be dominated by the *s*^2^ behavior. In this limit we have;

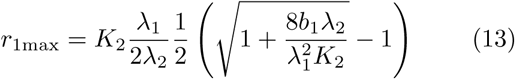

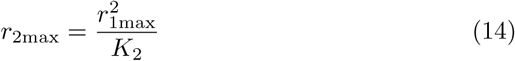

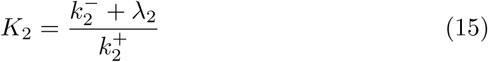

and, to leading order in *s*, we find;

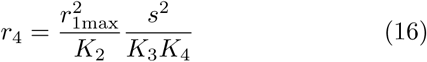

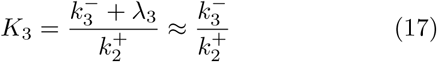

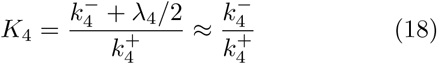

where *K*_2_, *K*_3_, and *K*_4_, are recognized as the dissociation constants for the processes (4)-(6). The asymptotes, Eq. (12) and Eq. (16), cross at the effective dissociation constant;

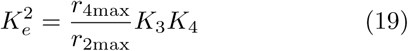

Assuming dimer forms are equally protected against proteolytic turnover, the effective dissociation constant will be larger than the dissociation constant for the second order binding of the ligand, (*K*_3_*K*_4_)^1*/*2^.

The effective independence, *ζ*_*e*_, is determined by setting the parametrization, Eq. (1), equal to the steady state value of *r*_4_ at *s*=*K*_*e*_. We find the independence term;

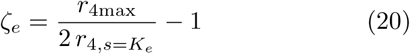

which has the value 1 for independent ligand bindings and 0 for fully cooperative bindings.

The timescale related to the changes in transcription factor concentration can be isolated by summing Eqs. (7)-(10). Each of the dimeric species contains two transcription factor constituents and is therefore counted twice. We then arrive at;

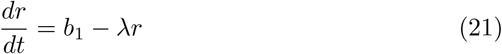

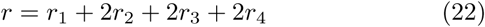

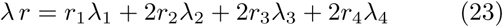

where *r* is the total transcription factor concentration and *λ* is the weighted average of the dilution and degradation rates. Eq. (21) governs the slow changes in the total transcription factor concentration that appear when the degradation rates differ.

### B. The independence measure

We will now try to gain some insight into why underlying independent ligand bindings do not necessarily lead to non-cooperative activation. The quasistatic approximations (fast on/off rates, 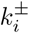, compared to turnovers, *λ*_*i*_) for Eqs. (5)-(6) lead to [18];

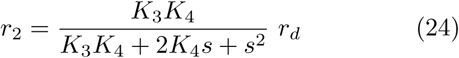

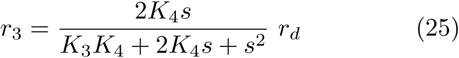

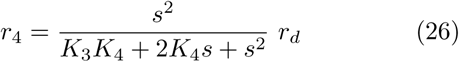

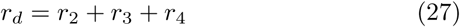

being satisfied at all times.

If *r*_*d*_ is constant, Eq. (26) is of the same form as Eq. (1) with effective dissociation constant *K*_*e*_ = (*K*_3_*K*_4_)^1*/*2^ and independence measure, 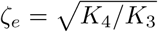. (Note that independent ligand bindings have *K*_3_ = *K*_4_ and results in the independence measure being 1.) However, if *r*_*d*_ depends on the ligand concentration, *s*, the independence in the underlying reaction need not be inherited by the overall reaction. We will now look into, when this is the case. For simplicity, let us assume that all dimer forms of the transcription factor are equally protected against proteolytic degradation, i. e. *λ*_2_ = *λ*_3_ = *λ*_4_ = *λ*_*d*_. In the static limit, the total dimer concentration may be written;

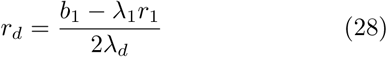

If the dimers are stable 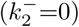, the numerator is a constant, and so is *r*_*d*_. The effective dissociation constant as well as independence in the underlying reaction is therefore inherited by the overall reaction.

However, if the dimers are unstable 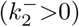, the dimer concentration in Eq. (28) becomes a function of the ligand concentration, *s*. In order for Eq. (1) to represent the coupled processes, we can remedy this by allowing for a modified effective dissociation constant and independence as described by Eqs. (19) and (20).

Similarly, if the active dimer has lower proteolytic degradation than the other dimer forms, *r*_*d*_ will depend on the ligand concentration and, again, this leads to modified effective independence.

### C. Cooperativity by binding to cognate operator (DNA) site

So far, we have investigated the production and activation of the transcription factor through ligand binding. Let us now consider what happens when the activated transcription factor binds to its cognate operator site on the DNA as indicated in Fig. 1. The concentration of operator sites on the DNA that is occupied by an activated transcription factor is denoted *p*_*a*_=[R_2_S_2_-*P*_*X*_] and the total concentration of operator sites is denoted *p*_*t*_=[*P*_*X*_] + [R_2_S_2_-*P*_*X*_]. The process is in quasistatic equilibrium (fast on/off rates) and therefore the concentration of occupied DNA sites is;

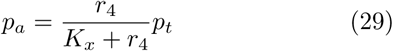

where 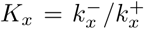 is the dissociation constant for the process. Insertion of *r*_4_ from Eq. (1) and rearranging results in;

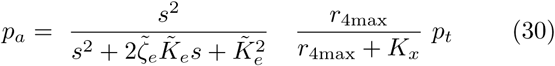

which is of the same functional form as the activated transcription factor concentration, but now with appropriately rescaled independence measure and dissociation constant;

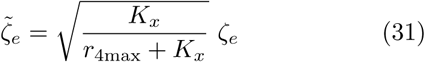

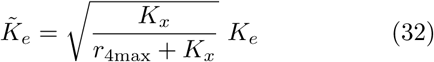

High transcription factor levels, *r*_4max_ ≫ *K*_*x*_, will push the activation towards cooperative behavior, i. e. 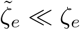. This criterion is easily satisfied with typical experimentally observed dissociation constants and transcription factor concentrations [19].

This way of generating cooperative activation generalizes to higher order activation. Furthermore, the activation in downstream first-order processes resulting from the activation will look increasingly cooperative for each step. In this sense, cooperative activation constitutes a stable fixed point as a functional form. We express this symbolically as;

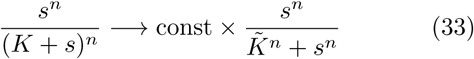

where the arrow signifies convergence for large activating transcription factor concentrations or higher number of first order steps in the sequence of processes. For example, even for 4^th^ order activation, essentially full cooperativity is easily established with a single activation followed by a first order process.

In the literature, the parametrization 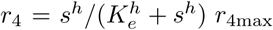 is frequently used as a proxy for the transcription factor-ligand concentration. However, it fails to simultaneously reproduce the asymptote at *s* ≪ *K*_*e*_ and the behavior around *s* ∼ *K*_*e*_. Further, it fails to capture the convergence towards cooperative binding as in Eqs. (31) and (33).

## RESULTS AND DISCUSSION

We have presented two mechanisms that make the sensing of a ligand through activation of transcription factors and subsequent binding to their cognate operator site on the DNA appear cooperative. The apparent cooperativity of the activation of the dimeric transcription factors was found to be possible in systems where dimerization drives ligand binding with unstable dimers. The apparent cooperativity generated when activated transcription factors bind to their cognate site on the DNA only requires the maximal transcription factor concentration to be higher than the dissociation constant for the binding to DNA.

Both mechanisms can be seen as a consequence of having two saturation processes on top of each other. When the dimer is unstable, the initial balance between the monomer form and the dimer form will be shifted towards the dimer form as the ligand concentration increases. This saturation process happens in parallel with the underlying shifting between the R_2_ form and ligand bound R_2_S and R_2_S_2_ forms of the transcription factor. As a result, the effective dissociation constant shifts to higher values than the underlying. Therefore, at the effective dissociation constant, the denominator in Eq. (26) has already shifted to *s*^2^ behavior. This leads to the observed cooperative behavior. With this clarification, we expect the mechanism to generalize to higher order activation of transcription factors. The explanation for the cooperative behavior as the active transcription factor binds on the DNA is similar, except, here the effective dissociation constant shifts to lower values, but again with the effect that the independence is quenched at ligand concentration approximating to the effective dissociation constant.

In Fig. 2, two examples with independent underlying ligand bindings are shown. As can be seen from the plots in the upper panels, both display cooperative behavior in the activation of the transcription factor. Figs. 2a-b are log-linear plots, stressing the behavior around *K*_*e*_. The log-log plots in Figs. 2c-d provide a representation of the full dynamic range of the activation and give a clear view of the asymptotic behavior. The parameters used in the calculations are shown in Table I (indicated as “cooperative”) and are all within the referenced physiological ranges. In the lower panels, the time response for the two “cooperative” parameter sets are shown. This demonstrates, that the system is structured such that through appropriate evolution of parameters it can effect both slow and fast responses to a sudden increase in the ligand concentration.

**TABLE I.** Parameters used in the model where dimerization drives ligand binding in Fig. 2. The two leftmost parameter sets lead to cooperative activation at the promoter as shown in Fig. 2. The “Low Pass” set has slow dimer dilution and fast monomer dilution which gives rise to a noise-reducing slow response in transcription factor activation, i. e. analogous to low-pass filters. The “All Pass” set has equal dimer and monomer dilution rates which lead to all-pass characteristics, i. e. a fast response. The dilution rates include proteolytic degradation as well as dilution by cell division. The two rightmost parameter sets lead to non-cooperative activation at the promoter, one with unstable dimer, the other with stable dimer.

| Par | Cooperative |  | Non-cooperative |  | Description | Ref. |
| --- | --- | --- | --- | --- | --- | --- |
| | Low Pass | All Pass | Unstable $R_2$ | Stable $R_2$ | | |
|  | Fig. 2, left | Fig. 2, right |  |  | Illustration |  |
| $b_1$ | 1000 nM/h | 1000 nM/h | 1000 nM/h | 1000 nM/h | Constitutive transcription factor production | [20, 21] |
| $k_2^+$ | $0.05 \text{ nM}^{-1}\text{h}^{-1}$ | $0.05 \text{ nM}^{-1}\text{h}^{-1}$ | $0.05 \text{ nM}^{-1}\text{h}^{-1}$ | $7.2 \text{ nM}^{-1}\text{h}^{-1}$ | $2R \rightarrow R_2$ rate constant | [21–23] |
| $k_2^-$ | $100 \text{ h}^{-1}$ | $4000 \text{ h}^{-1}$ | $100 \text{ h}^{-1}$ | $0 \text{ h}^{-1}$ | $2R \leftarrow R_2$ rate | [19, 21] |
| $k_3^+$ | $100 \text{ nM}^{-1}\text{h}^{-1}$ | $100 \text{ nM}^{-1}\text{h}^{-1}$ | $10 \text{ nM}^{-1}\text{h}^{-1}$ | $10 \text{ nM}^{-1}\text{h}^{-1}$ | $R_2 + S \rightarrow R_2S$ rate constant | [21] |
| $k_3^-$ | $1000 \text{ h}^{-1}$ | $1000 \text{ h}^{-1}$ | $1000 \text{ h}^{-1}$ | $1000 \text{ h}^{-1}$ | $R_2 + S \leftarrow R_2S$ rate | [21] |
| $k_4^+$ | $100 \text{ nM}^{-1}\text{h}^{-1}$ | $100 \text{ nM}^{-1}\text{h}^{-1}$ | $10 \text{ nM}^{-1}\text{h}^{-1}$ | $10 \text{ nM}^{-1}\text{h}^{-1}$ | $R_2S + S \rightarrow R_2S_2$ rate constant | [21] |
| $k_4^-$ | $1000 \text{ h}^{-1}$ | $1000 \text{ h}^{-1}$ | $1000 \text{ h}^{-1}$ | $1000 \text{ h}^{-1}$ | $R_2S + S \leftarrow R_2S_2$ rate | [21] |
| $\lambda_1$ | $20 \text{ h}^{-1}$ | $10 \text{ h}^{-1}$ | $0.5 \text{ h}^{-1}$ | $20 \text{ h}^{-1}$ | Monomer dilution rate | [9, 10] |
| $\lambda_d$ | $0.5 \text{ h}^{-1}$ | $10 \text{ h}^{-1}$ | $0.5 \text{ h}^{-1}$ | $0.5 \text{ h}^{-1}$ | Dimer dilution rate | [11, 13, 14] |
| $r_{4\text{max}}$ | 172 nM | 19.1 nM | 951 nM | 847 nM | Maximal dimer concentration | |
| $(K_3K_4)^{1/2}$ | 10 nM | 10 nM | 100 nM | 100 nM | Underlying dissociation constant | |
| $(K_4/K_3)^{1/2}$ | 1 | 1 | 1 | 1 | Underlying independence | |
| $K_e$ | 120 nM | 125 nM | 138 nM | 100 nM | Effective dissociation constant | |
| $\zeta_e$ | 0.09 | 0.15 | 0.91 | 1.00 | Effective independence | |

**FIG. 2.**
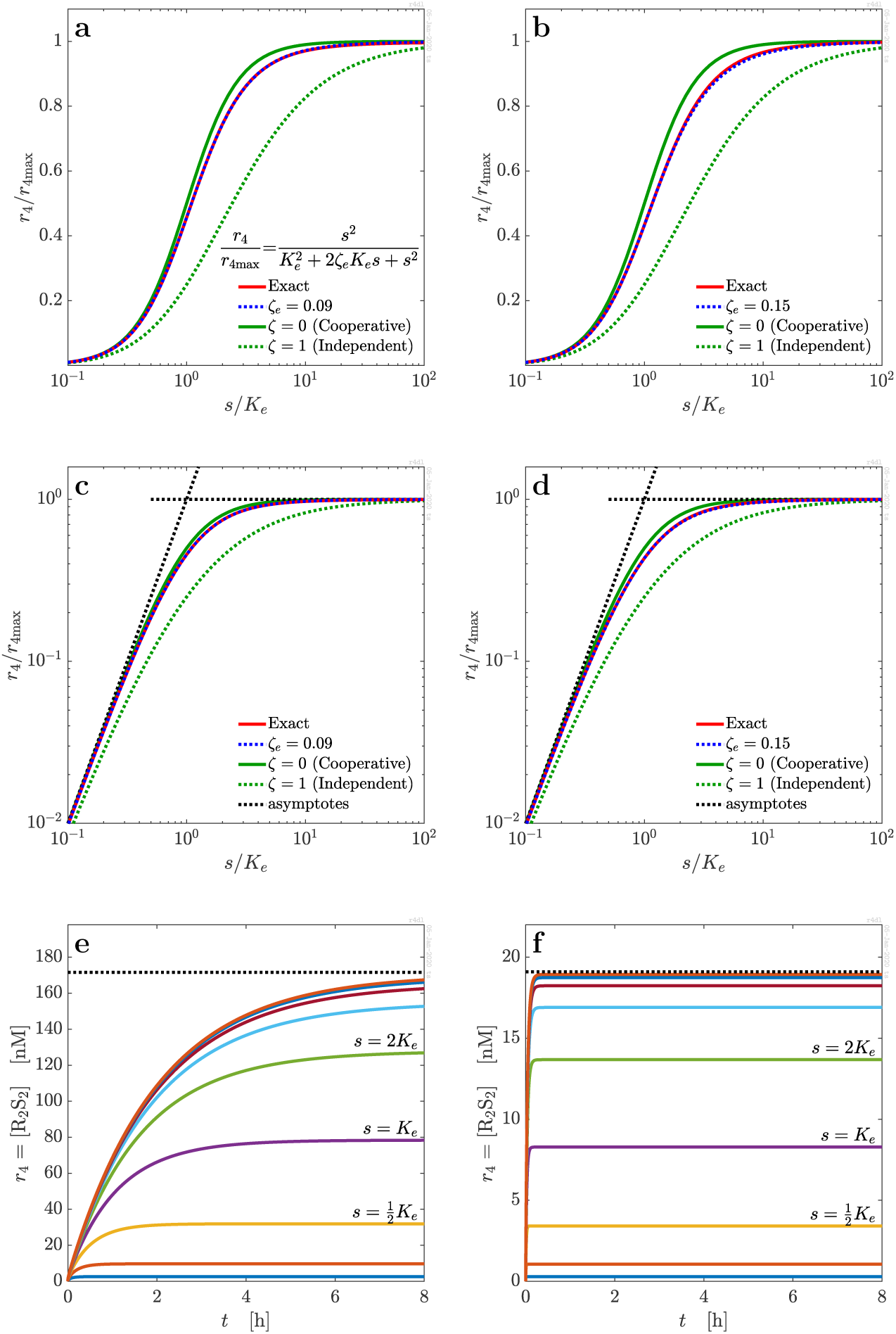
Cooperative activation of dimeric transcription factor. Activated transcription factor concentration as a function of ligand concentration for the model where dimerization drives ligand binding with parameter sets from Table I. Semi-log (panels a and b) as well as log-log (panels c and d) versions are shown. Slow response is shown in panels a, c, e. Fast response is shown in panels b, d, f. Panels a, c and b, d: We observe that transcription factor activation appears almost fully cooperative even though the underlying ligand bindings are independent. Panels c) and d): Time response following introduction of the ligand at concentrations *s* = 2^*n*^*K*_*e*_, *n* = −3, −2, …, 5.

In Table I, we also show two examples which display non-cooperative behavior in the activation of the transcription factor. They follow the behavior of the plots with effective independence (*ζ*_*e*_ = 1) in Fig. 2. The left column has unstable dimers, whereas the right column has stable dimers, thereby covering typical situations. Both parameter sets lead to a fast response.

We note that it is possible to construct parameter sets with stable dimers and cooperative transcription factor activation, though, with less typical proteolytic degradation rates (*λ*_2_ ≫ *λ*_3_ = *λ*_4_).

The cooperativity and the filtering properties of the transcription factor activation depend on the basal production of transcription factors. Substituting the promoter driving expression of the transcription factor by a heterologous promoter with stronger or weaker transcription rate will therefore result in altered cooperativity and filtering properties. Similarly, increasing the maximal transcription factor concentration, e. g., by expressing it from a multi-copy plasmid, will modify the overall cooperativity, the effective dissociation constant, and the response time of the regulatory motif.

We have given a couple of typical examples of regulatory systems which display cooperative behavior without requiring cooperativity at the elementary level. We expect that simple mechanisms similar to those presented here may apply to other regulatory systems as well.

## III. CONCLUSION

The primary finding from our analysis is that the cooperative activation of transcription factors can be established even with *independent* underlying ligand bindings. The binding of the activated transcription factor complex to its cognate operator site on the DNA pushes the regulation even further in the direction of cooperative behavior. Our analysis accounts for the inherent cooperativity in the biological activity of many dimeric transcription factors.

A number of advantages are associated with the cooperative activation of transcription factors, and most are related to improvements of basic sensory function. Cooperative activation can provide an expansion of the dynamic range of gene expression, and it can improve stability when occurring in feed-back systems that control rapid switching between different responses. It is therefore important that we have modelling approaches that can account for cooperativity in sensory systems where underlying cooperative mechanisms may not be present, thus, explaining cooperativity without the need of elaborate evolutionary constraints.

## Appendix: Ligand binding drives dimerization

For completeness, we include the simplest version of a regulatory motif where ligand binding drives dimerization

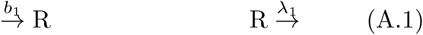

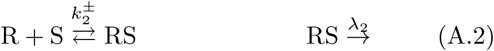

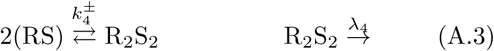

The corresponding kinetic equations are

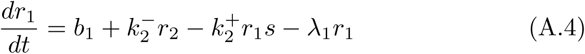

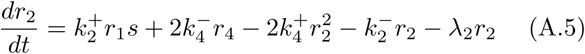

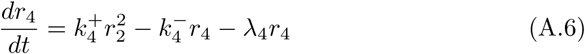

with combinatorial weights explicitly included. We employ the convention, that reactions are numbered by the number of constituents in the resulting product.

Again, in the static limit of the activation, the form Eq. (1) provides an excellent description. To determine *K*_*e*_ we need the asymptotic behavior at small *s* and at large *s* in the static limit of the kinetic equations. The derivation is analogous to the derivation of Eqs. (11)-(19). With dissociation constants defined as;

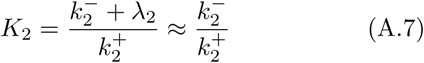

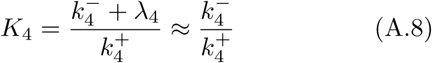

the asymptotic behavior at small *s* is given by;

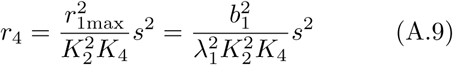

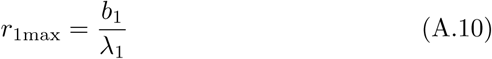

The maximal RS and activated transcription factor levels are reached at large *s*;

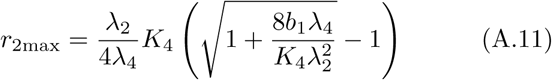

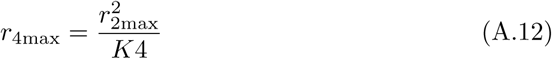

The effective dissociation constant sits at the crossing of the asymptotes;

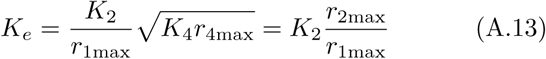

and is typically different from *K*_2_.

In the limit of low- or high-level production of transcription factors we find;

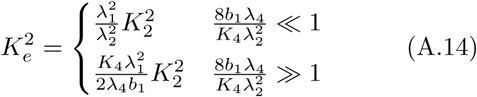

In the case where the production of transcription factors is low and the monomer forms have similar degradation rates, this does not lead to a modification of the dissociation constant. However, when there is large production of transcription factors, the rescaling of the dissociation constant can be significant. In the case where ligand binding drives dimerization, we did not find cooperativity in the activation with physiological values of the parameters that we tested. However, the transcriptional activation at the promoter may still be cooperative as a consequence of the mechanism described in “Cooperativity by binding to cognate operator (DNA) site”.

## References

[1] F. Crick, Nature 227, 561 (1970).

[2] D. S. Latchman, The International Journal of Biochemistry & Cell Biology 29, 1305 (1997).

[3] C. Fuqua, S. C. Winans, and E. P. Greenberg, Annual Review of Microbiology 50, 727 (1996).

[4] P. Williams, Expert Opin. Ther. Targets 6, 257 (2000).

[5] A. Eberhard, J. Bacteriology 109, 1101 (1972).

[6] K. H. Nealson, T. Platt, and J. W. Hastings, J. Bacteriol. 104, 313 (1970).

[7] S. H. Choi and E. P. Greenberg, J. Bacteriology 174, 4064 (1992).

[8] J. Ferkinghoff-Borg and T. Sams, Mol. BioSyst. 10, 103 (2014).

[9] J. Zhu and S. C. Winans, Proc. Natl. Acad. Sci. USA 96, 4832 (1999).

[10] J. Zhu and S. C. Winans, Proc. Natl. Acad. Sci. USA 98, 1507 (2001).

[11] J. E. Gonzalez and N. D. Keshavan, Microbiology & Molecular Biology Reviews 70, 1 (2006).

[12] M. J. Bottomley, E. Muraglia, R. Bazzo, and A. Carfi, Journal of Biological Chemistry 282, 13592 (2007).

[13] K. J. Sappington, A. A. Dandekar, K.-I. Oinuma, and E. P. Greenberg, mBio 2, e00011 (2011).

[14] A. Claussen, T. H. Jakobsen, T. Bjarnsholt, M. Givskov, M. Welch, J. Ferkinghoff-Borg, and T. Sams, International Journal of Molecular Sciences 14, 13360 (2013).

[15] A. V. Hill, Proceedings of the Physiological Society, iv (1910).

[16] C. Bohr, Zentralblatt für Physiologie XVIII, 682 (1904).

[17] C. Bohr, K. Hasselbalch, and A. Krogh, Skandinavisches Archiv Für Physiologie 16, 402 (1904).

[18] M. Welch, J. Gross, J. T. Hodgkinson, D. R. Spring, and T. Sams, Biochemistry 52, 4433 (2013).

[19] K. Sneppen, Models of Life (Cambridge University Press, Cambridge, 2014).

[20] M. Fagerlind, S. Rice, P. Nilsson, M. Harlen, S. James, T. Charlton, and S. Kjelleberg, Journal of Molecular Microbiology and Biotechnology 6, 88 (2003).

[21] M. G. Fagerlind, P. Nilsson, M. Harlén, S. Karlsson, S. A. Rice, and S. Kjelleberg, BioSystems 80, 201 (2005).

[22] M. Schlosshauer and D. Baker, Protein Sci 13, 1660 (2004), 0131660[PII], 15133165[pmid].

[23] M. Schlosshauer and D. Baker, J. Phys. Chem. B 106, 12079 (2002).

